# Enhanced processing of aversive stimuli on embodied artificial limbs by the human amygdala

**DOI:** 10.1101/2021.06.14.448367

**Authors:** Antonin Fourcade, Timo Torsten Schmidt, Till Nierhaus, Felix Blankenburg

## Abstract

Body perception has been extensively investigated, with one particular focus being the integration of vision and touch within a neuronal body representation. Previous studies have implicated a distributed network comprising the extrastriate body area (EBA), posterior parietal cortex (PPC) and ventral premotor cortex (PMv) during illusory self-attribution of a rubber hand. Here, we set up an fMRI paradigm in virtual reality (VR) to study whether and how threatening (artificial) body parts affects their self-attribution. Participants (N=30) saw a spider (aversive stimulus) or a toy-car (neutral stimulus) moving along a 3D-rendered virtual forearm positioned like their real forearm, while tactile stimulation was applied on the real arm in the same (congruent) or opposite (incongruent) direction. We found that the PPC was more activated during congruent stimulation; higher visual areas and the anterior insula (aIns) showed increased activation during aversive stimulus presentation; and the amygdala was more strongly activated for aversive stimuli when there was stronger multisensory integration of body-related information (interaction of aversiveness and congruency). Together, these findings suggest an enhanced processing of aversive stimuli within the amygdala when they represent a bodily threat.

## 1 Introduction

In an iconic scene of the classic James Bond movie Dr. No (1962), the spy is shown taking a well-deserved rest in his bed when he suddenly feels a surprising touch on his upper arm. He slowly turns his head toward the source of this unexpected sensation and sees a tarantula crawling onto his shoulder, and as the fear sets in, beads of sweat form on his forehead as the spider crawls towards his head. This typical fear response arises, because Bond perceives that the spider poses a bodily threat to him. How does he infer that what he sees, the spider, is also stimulating what he feels on his arm? This is a classic example of multisensory integration, i.e., stimuli registered by distinct modalities (e.g., sight and touch) are inferred to be caused by the same source, i.e., the spider on the skin ^1–3^. These inference and integration processes are highly plastic, and research on body ownership has explored how even body representations can be influenced and adjusted depending on incoming (multi-)sensory information ^4,5^ using the Rubber Hand Illusion ^6,7^ (RHI). The RHI is induced when the rubber hand and the participant’s real hand (hidden from sight) are stroked in a temporally and spatially congruent manner ^8,9^. The visual and tactile information are integrated in the brain, while the incoming proprioceptive information from the real hand is seemingly down-weighted. At the same time, participants may experience the illusion of perceiving the rubber hand as part of their body. Functional neuroimaging studies using the RHI have revealed a brain network ^10,11^, comprising the body-selective extrastriate body area (EBA), posterior parietal cortex (PPC), and ventral premotor cortex (PMv), which is thought to integrate sensory information in order to recalibrate peripersonal space ^12–14^, to support action ^15,16^. The RHI paradigm has also been used to investigate how emotion processing, related to threat, interacts with the illusionary self-attribution of the fake hand. Ehrsson and colleagues (2007) showed that the anterior insula (aIns) and the anterior cingulate cortex (ACC), which are part of an interoceptive network implicated in physiological and emotional processing ^17,18^, were activated when the rubber hand was threatened with a needle while participants experienced the RHI, and that activation in these regions was correlated with participants’ subjective ratings of ownership. The researchers posited that the involvement of interoceptive brain regions during the RHI may add to the vividness of the body ownership experience by including homeostatic emotional components (e.g., pain anticipation, temperature) ^19^. However, it is yet not well understood how the multisensory integration underlying the sense of body ownership may mediate emotion - especially at an early phase of this integration, before the full onset of the actual illusion is experienced ^8^. Another key brain region involved in emotional and sensory processing is the amygdala, known for its role in processing fear and emotional salience of external stimuli ^20^. Peelen and colleagues (2007) reported that the EBA showed greater activation when participants viewed body movements representing basic emotions (e.g., bodily expressions of fear or anger) compared to neutral body movements (e.g., walking, jumping), and that the activation of the amygdala was correlated with this modulation of EBA activity ^21^, implicating the amygdala in body-related emotional processing. However, whether activation of the amygdala is modulated by the self-relevance of body parts – i.e., their self-attribution or “embodiment” - in the presence of an aversive stimulus remains an open question. And more generally, it is unclear how the brain processes emotionally loaded visuo-tactile stimuli that are related to a self-attributed body part, as compared to a non-self-attributed one.

Recently, virtual reality (VR) has emerged as a new tool to advance the investigation of both body ownership and emotion processing. The brain’s flexibility in representing body ownership has been emphasized by studies using the RHI paradigm in VR ^9^, as well as studies investigating a whole-body transfer illusion ^22,23^. VR also provides greater ecological validity ^24,25^; researchers are able to create more contextualized and realistic experiences, while still maintaining experimental control. Indeed, experiments using VR have shown that the elicitation of emotions is stronger when participants were more immersed in virtual environments ^26–28^. Following this, stereoscopic rendering via MRI-compatible VR goggles has been shown to be more immersive than the presentation of 2D stimuli ^29^ and a recent study on emotion regulation that combined VR and functional neuroimaging found activation of the amygdala when participants were immersed in a virtual environment combined with music designed to elicit anguish ^30^. Altogether, VR gives a unique opportunity to investigate the early phase of the multisensory integration underlying the RHI in the presence of emotional stimuli, while being able to record brain activity with fMRI.

Here we intended to investigate the relevance of emotional processing in the context of bodily threat by focusing on the neuronal correlates of processing aversive vs neutral stimuli on embodied (artificial) limbs. Specifically, we aimed our VR paradigm at extending the available methods of investigating multimodal integration in a controlled manner ^9,22,23^ while also adding an affective component to the stimulus: aversiveness ^31^. We designed an experiment in which we manipulated the congruency of visuo-tactile stimulation (congruent vs. incongruent) on a participant’s arm and the aversiveness of the stimuli (aversive versus neutral). Based on the RHI literature, we hypothesized that a similar network of regions, comprising the EBA, PCC, and PMv, will be activated for the integration of visual and tactile information. Secondly, considering the literature on emotion processing, we speculated that the aIns, ACC, and amygdala would show increased activation during the presentation of an aversive stimulus, as compared to a neutral one. Finally, we postulated that the amygdala, the aIns and the ACC would show an interaction effect, such that congruent-aversive stimulation will elicit higher activations than other stimulus pairings.

## 2 Methods

Participants underwent an fMRI scanning session followed by a retrospective questionnaire on their subjective body ownership experience and their emotional response to the stimuli. During the scan, participants were presented with visual objects moving either in the proximal-distal direction (i.e., from elbow to wrist) or distal-proximal direction (i.e., from wrist to elbow) on a virtual 3D-rendered forearm in the same position as their real arm. At the same time, tactile electric stimulation was applied moving in either the same direction as the visual stimulus or in the opposite direction, resulting in visuo-tactile congruent and incongruent trials, respectively. The moving objects were either aversive stimuli (spider) or neutral objects (toy car). Thus, the experimental design comprised a 2 × 2 × 2 factorial design with congruency, aversiveness, and direction of visual motion (left and right) as independent factors.

### 2.1 Participants

Thirty-three healthy participants (age range: 19–36 years; 20 females; all right-handed [mean Laterality Quotient = 90 as assessed with the Edinburgh Handedness Inventory ^32^]; normal or corrected-to-normal vision; no self-reported arachnophobia) participated in the experiment. Three participants’ datasets were excluded due to inattentiveness during the control task (see below), resulting in *N* = 30 datasets used for the analysis. All participants gave written informed consent before the experiment. The study was approved by the local Ethical Committee of Freie Universität Berlin and conducted in accordance with this approval and the relevant guidelines and regulations.

### 2.2 Experimental set-up

The participant’s right arm was placed horizontally across the chest using pillows for support, in a position corresponding to the presentation of the virtual arm. To ensure that the location of visual stimuli in eye-centered coordinates remained the same, participants were instructed to fixate a small red dot in the middle of the virtual forearm (and center of the virtual field of view) throughout the whole experiment. For full, direct vision of the virtual arm, the participant’s head was slightly tilted down towards the chest within the head coil (approx. 20-30°), and the head and neck were supported with foam padding. Stereoscopic goggles were attached both to the participant’s forehead and to the head coil with Velcro strips to minimize motion during the experiment. The participant’s real arm was completely occluded from view by the goggles. A fiber optic response button box (fORP, Current Designs, Philadelphia, PA) was placed in the left hand to collect responses to an attention task (see below).

#### 2.2.1 Paradigm

There was a total of eight trial types (Congruent vs. Incongruent × Aversive vs. Neutral × Proximal→ Distal vs. Distal→Proximal visual motion). Within each of the six runs, each condition was presented six times (i.e., 48 trials per run). Additionally, one attentional control trial, where the fixation dot briefly blinked (50-ms on/off period), was presented per condition and run (i.e., 8 trials per run); participants were instructed to respond to the blinking fixation dot with a button press. This resulted in a total of 56 trials per run and 336 trials overall. Trials were randomized within each run. Each trial lasted for two seconds, followed by a jittered inter-trial interval of two to six seconds (approx. 7-min per run). After the scan, participants were asked to fill out a questionnaire assessing participants’ subjective ratings of the congruency and aversiveness of the stimuli.

#### 2.2.2 Virtual Reality

Digital stereoscopic goggles (VisuaSTIM, 800×600 pixels, 30° eye field) and PsychToolbox 3.0.14 ^33,34^ with MATLAB 2016a 64bit (Mathworks, Massachusetts) were used to present a photorealistic 3D-rendered virtual arm in a plausible posture with respect to the real arm (i.e., an anatomically plausible configuration and location in space), with the hand palm down and in a fist (see Figure 1B).

**Figure 1.**
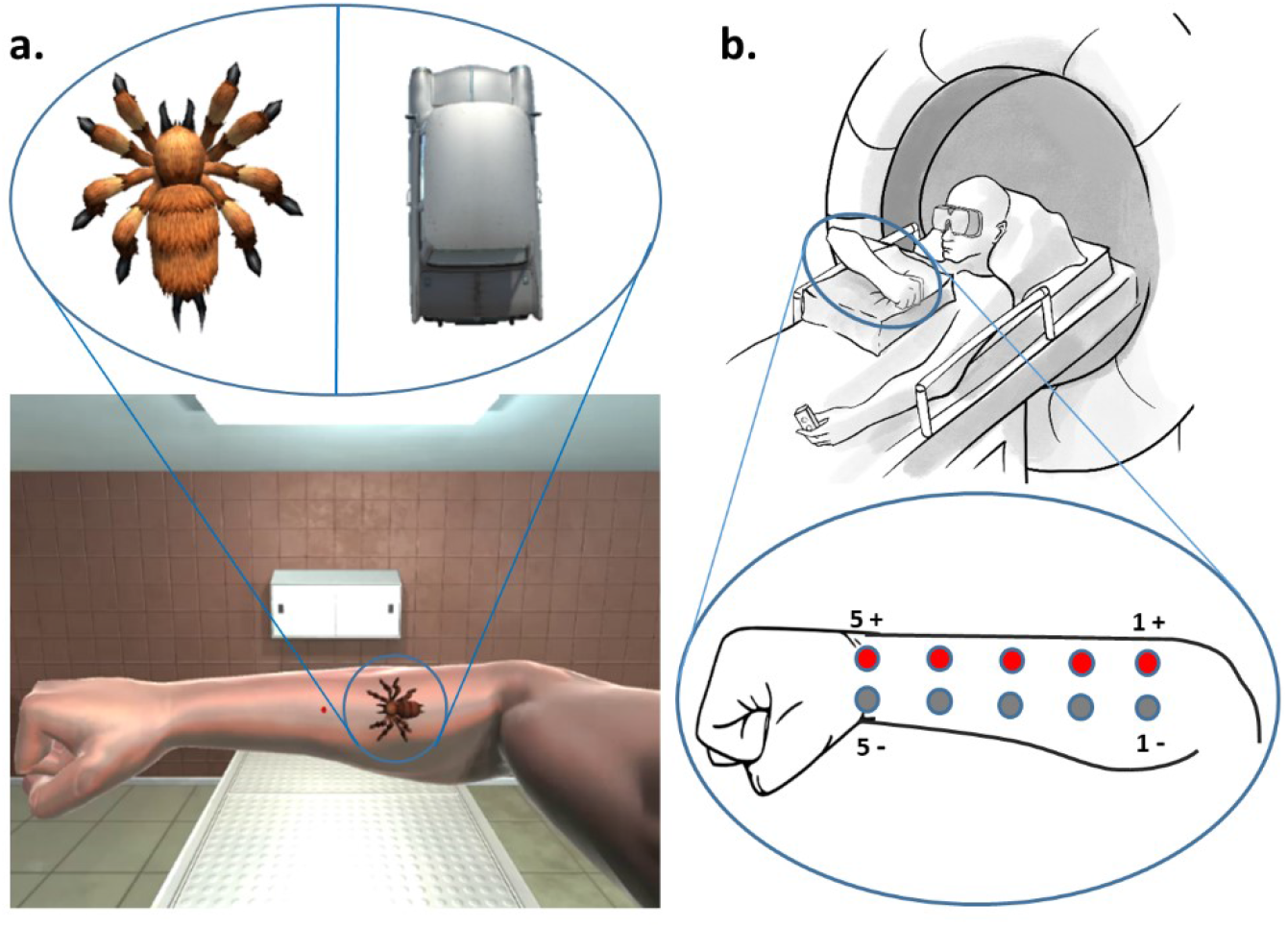
**a**. Image of the virtual environment seen by the participants. Only the right-eye view is shown. Stimuli used for aversive (spider) and neutral (car) conditions. The red fixation dot at the center of the forearm blinked briefly during attentional control trials. **b**. Details of the set-up inside the MRI room. Five pairs of surface-adhesive electrodes were positioned on the lateral side of the right forearm, from the wrist to the elbow, to enable five stimulation sites. The right arm was placed horizontally on top of the participant’s chest using pillows, fist closed. The left hand was holding the response button box with the arm along the body. The head was slightly tilted in the direction of the chest within the head coil (approx. 20-30°). Stereoscopic goggles were attached to the participant’s forehead and the head coil (not shown) with Velcro strips to minimize motion.

The stimulus presentation computer was equipped with a NVidia GeForce GTX 750Ti graphic card with two display outputs (one for each eye). For each condition, stereoscopic videos were created with the Unity3D 2017 software package (Unity Technologies, California) and 3D assets available on the Unity Store (https://assetstore.unity.com). The aversive and neutral stimuli were designed to be as similar as possible, e.g. with the same size, moving speed and starting/end point. However, the legs of the spider were moving to simulate crawling, while the shape of the car was not changing. The background of the videos consisted of a neutral room with a bench, a cabinet and a ceiling light, as well as the virtual right arm with a red dot in the middle (see Figure 1A). In Unity3D, the distance between the two recording cameras simulating both eyes (and generating the stereoscopic videos) was set to the mean adult interpupillary distance of 63mm ^35^ to create a 3D effect.

#### 2.2.3 Electrostimulation

Prior to the scanning, five pairs of surface-adhesive electrodes were positioned on the lateral side of the right forearm, from the wrist to the elbow (see Figure 1B). A constant current neurostimulator (DS7A, Digitimer, Hertfordshire, United Kingdom) was used to deliver electrical pulses (square wave, 0.2 ms duration, mean intensity = 5.94 ± 1.46mA) to the five stimulation sites. During the experiment, the same stimulus intensity was used for all sites. Electrode positions were adjusted individually so that the pulses from each electrode had comparable intensity and could be spatially discriminated, without producing discomfort, radial stimulation, or muscle contractions. An 8-channel relay card (RX08-LPT, GWR Elektronik) was used to control the administration of pulses. The relay card was operated with MATLAB via the parallel port (LPT) of the computer. Five pulses were always delivered sequentially (500-ms delay) and could start at either the left (wrist) or right (elbow) electrode. The intensity was adjusted to each participant such that they reported a tingling sensation that resembled an insect crawling on their arm.

#### 2.2.4 fMRI data acquisition

The experiment was conducted using a 3 Tesla scanner (Tim Trio, Siemens, Germany) equipped with a 12-channel head coil. T2*-weighted images were acquired using a gradient echo-planar imaging sequence (3 × 3 × 3 mm voxels, 20% gap, matrix size = 64 × 64, TR = 2000ms, TE = 30ms, flip angle = 70°). Six runs with 176 functional volumes each were recorded for each participant. After the functional runs, a gradient-echo (GRE) field map (3 × 3 × 3 mm voxels, TR = 488ms, TE1 = 4.92ms, TE2 = 7.38ms, 20% gap, flip angle = 60°), and a high-resolution T1-weighted structural image was acquired for each participant (3D MPRAGE, voxel size = 1 × 1 × 1 mm, FOV = 256 × 256 mm, 176 slices, TR = 1900ms, TE = 2.52ms, flip angle = 9°).

#### 2.2.5 Data preprocessing and analysis

Data were processed and analyzed using SPM12 (Welcome Department of Cognitive Neurology, London, UK: www.fil.ion.ucl.ac.uk/spm/). Images were realigned to the first image of each run to correct for head motion. Each participant’s structural image was co-registered with the realigned functional images, and segmented into white matter (WM), gray matter (GM), and cerebrospinal fluid (CSF). Functional images were spatially normalized to the MNI space using DARTEL ^36^ and spatially smoothed by an isotropic Gaussian kernel of 8 mm full width at half maximum. Data were detrended using a linear mean global signal removal script ^33^. To reduce physiological and systemic noise in the functional data, the first five principal components accounting for the most variance in the CSF and WM signal time-courses, respectively, and the six realignment parameters, were added to the first-level general linear models (GLMs) as regressors of no interest ^37^. Each trial type was modeled as a regressor with a boxcar function (2-s duration) and convoluted with the standard hemodynamic response function from SPM. Attentional control trials were not included.

#### 2.2.6 Behavioral data

During the scan, button presses were recorded and d’ was calculated as an index of maintained attention. Hits were defined as a button response during an attentional control trial; false alarms as a button response during test trials. Perfect rates (p_hits_ = 1 or p_false alarms_ = 0) were corrected according to the 1/2N rule ^38,39^.

The post-scan questionnaire consisted in 15-items assessing participants’ subjective ratings of the congruency and aversiveness of the stimuli. To validate that participants perceived the visual and tactile stimuli as synchronous, they were asked “Was the visual moving object synchronized with the tactile stimulation?” The questionnaire also inquired whether participants were aware there was congruent and incongruent visual-tactile stimulation (“Was the tactile stimulation for some trials going the same/opposite direction as the visual moving object?”; one question per congruency condition). To assess the degree to which participants might have experienced the “ownership illusion,” they were asked to rate the following statements (based on ^6,7^), for congruent and incongruent visual-tactile stimulation separately: “I felt as if I was looking at my own arm and hand;” “I felt as if the virtual arm and hand was part of my body;” “I felt as if the virtual arm and hand were my arm and hand. Body Ownership scores for congruent and incongruent trials were then calculated separately by averaging the ratings. Finally, aversiveness of the spider and car were assessed by asking “Did the moving object make you feel uncomfortable/scared/pleased?” Each item was rated on a seven-point numerical rating scale ranging from “not at all” (0) to “definitely yes” (6).

#### 2.2.7 Statistical Analyses

For the fMRI data, on the first level, four trial conditions (Congruency × Aversiveness; left and right motion direction pooled together) were modeled as regressors, as well as four first-order time-modulated regressors. In addition, five CSF/WM components and the six motion parameters were added, resulting in 19 regressors. Four t-contrasts corresponding to the trial conditions were computed for each participant. On the second level, the first-level con-images were used to perform a 2 × 2 ANOVA with Congruency (Congruent, Incongruent) and Aversiveness (Aversive, Neutral) as factors.

Predefined ROI masks for the bilateral amygdala, insula, and ACC were created using the SPM Anatomy toolbox v3.0 ^40^. Because no atlas includes a map specifically for EBA and/or PMv, a 10 mm radius spherical ROI was created, centered on coordinates reported in an independent study ^11^ (left EBA, x = –50, y = –74, z = 6; right EBA, x = 54, y = –68, z = 2; left PMv, x = –52, y = 8, z = 28; right PMv, x = 52, y = 10, z = 32). This study was chosen because it was one of the first showing the involvement of EBA in the RHI and the reported coordinates are also consistent with later studies (e.g., ^41^).

We then performed a whole-brain analysis with family-wise error correction (FWE) at the cluster level (*p* < .05) using an initial voxel-wise threshold of *p* < .001, uncorrected. Then, following our *a priori* hypotheses, we additionally report results at *p* < .001, uncorrected within predefined regions of interest, i.e., left/right EBA, left/right PMv, left/right aIns, left/right ACC and left/right amygdala.

For post-hoc tests, we extracted the contrast estimates at the peak activation voxel for each pair of conditions and each subject. Pairwise t-tests were then performed, correcting for multiple comparisons using Bonferroni correction.

To check for potential effect of habituation of amygdala activity in response to aversive stimuli, we carried out an additional analysis, a 2 × 2 ANOVA with this time the first-order time-modulated regressors at the first level. We computed a negative contrast on the aversive congruent and incongruent parametric regressors, in order to investigate a time parametric modulation in the aversive trials.

Regarding the attention task, a one-way ANOVA was performed with Run as a factor with six levels to confirm continuous attentiveness to the task.

All ratings collected with the post-scan questionnaire were tested for normality with Shapiro-Wilk tests. As they did not pass these tests for normality, they were analyzed using non-parametric Wilcoxon’s signed-rank tests with *α* = 0.05. *P*-values concerning the stimulus affective ratings (“uncomfortable”, “scared’’, and “pleased”) were corrected for false discovery rate (FDR).

#### 2.2.8 Control Analyses

After the data collection, we aimed to investigate the potential effects of different individual levels of fear of spiders in our sample. The Fear of Spiders Questionnaire ^42^ (FSQ) was administered retrospectively via email. A subset of 25 participants responded (FSQ scores mean 17.4 ± 18.91). We carried out two additional control analyses on this subset of data. First, we performed a 2 × 2 ANOVA with Congruency (congruent, incongruent) and Aversiveness (aversive, neutral) as factors, in which the individual FSQ scores were added as a covariate of no interest. Second, we split participants into two groups (based on ^43^): low (FSQ scores < 15; n = 15) and high (n = 10) fear. We then performed a 2 × 2 ANOVA with Aversive-Congruency (aversive-congruent, aversive-incongruent) and Fear Group (low, high) as factors.

## 3 Results

### 3.1 Behavioral results

Participants’ attention, as indexed by d’, ranged from 0 to 3.07 per run. Across participants, d’ did not significantly differ between runs, *F*(5,179) = 0.18, *p* = .27. Each participant’s mean d’ across runs was calculated and an exclusion criterion of mean d’ = 1.66 was set. Three participants were excluded due to poor performance on the attention task.

Participants reported that they were able to identify that there were congruent trials (mean = 5.70, STD = 0.99), *Z* = 5.12, *p* < .001, and incongruent trials (mean = 5.73, STD = 0.74), *Z* = 5.15, *p* < .001. Tactile stimulation subjectively synchronized with the movement of the objects (mean = 5.33, STD = 1.18), *Z* = 4.73, *p* < .001. Body ownership ratings were higher for congruent stimulation (mean = 3.12, STD = 1.62) than incongruent stimulation (mean = 2.40, STD = 1.69), *Z* = 3.27, *p* < .001. P-values concerning the stimulus aversiveness ratings were corrected for false discovery rate (FDR); ratings for “uncomfortable” were significantly higher for spider (mean = 2.30, STD = 2.31) than for car (mean = 0.60, STD = 1.07), *Z* = 3.72, *p* < .001. Ratings for “scared” were higher for spider (mean = 1.8, STD = 1.91) than for car (mean = 0.3, STD = 0.79), *Z* = 3.51, *p* < .001. Ratings for “pleased” were significantly lower for spider (mean = 0.8, STD = 1.26) than for car (mean = 1.6, STD = 1.81), *Z* = 2.56, *p* = .009 (see also Figure S1 and Table S2 in Supplementary Material).

### 3.2 fMRI results

#### 3.2.1 Congruence vs. Incongruence

Contrasting congruent versus incongruent stimulation revealed higher activation within the PPC for congruent compared to incongruent trials. The left superior parietal lobule SPL (area 7A; see Figure 2A and Table 1) showed higher activation in the whole-brain analysis with FWE-correction. When testing in the a priori defined ROIs, the PMv and EBA showed no significant difference in activation between trials at a significance threshold of p < .001.

**Table 1.**
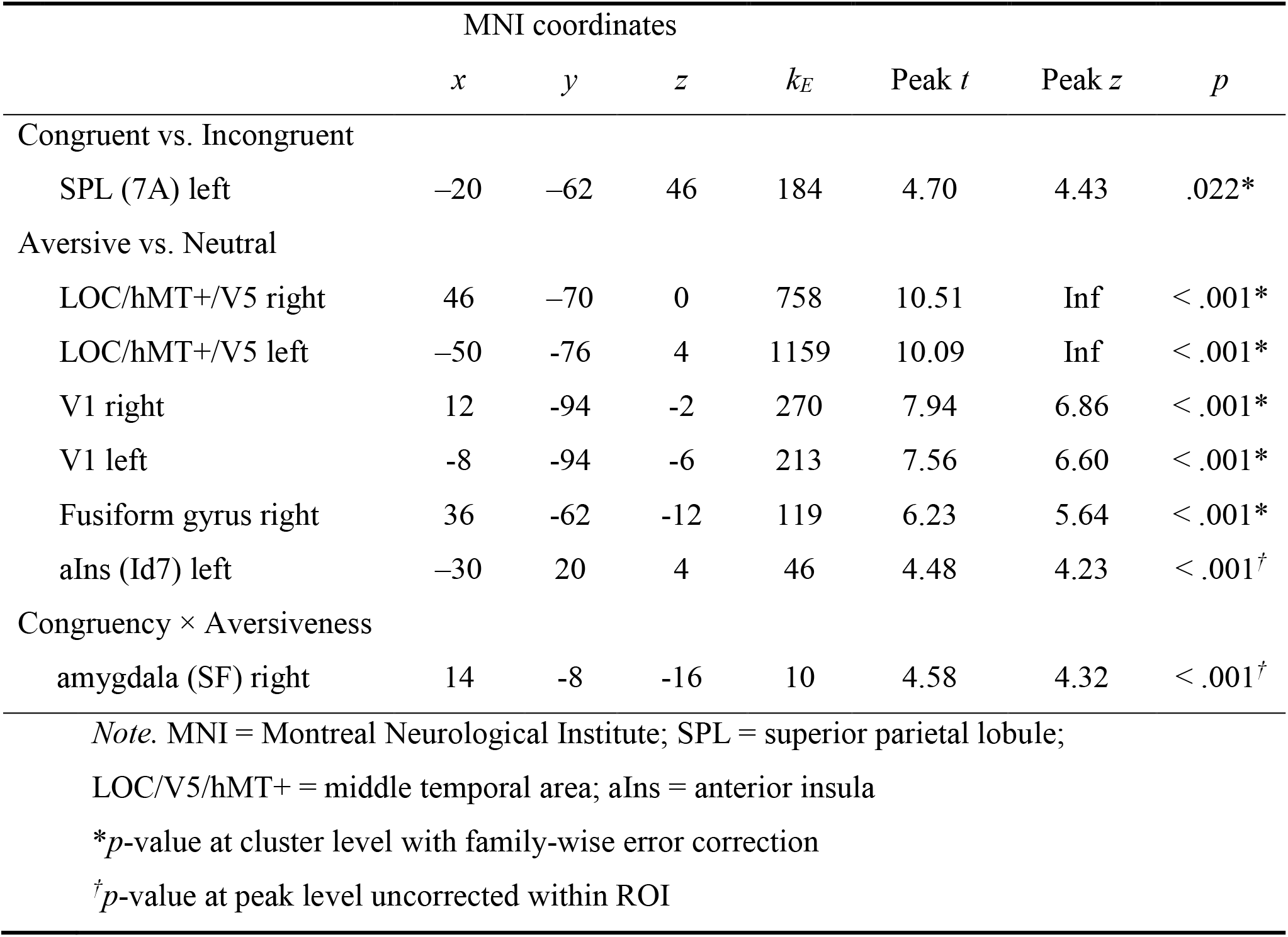
Significant activation differences obtained from contrasting congruent versus incongruent visual-tactile stimulation, aversive versus neutral conditions, and the interaction between congruency and aversiveness.

**Figure 2.**
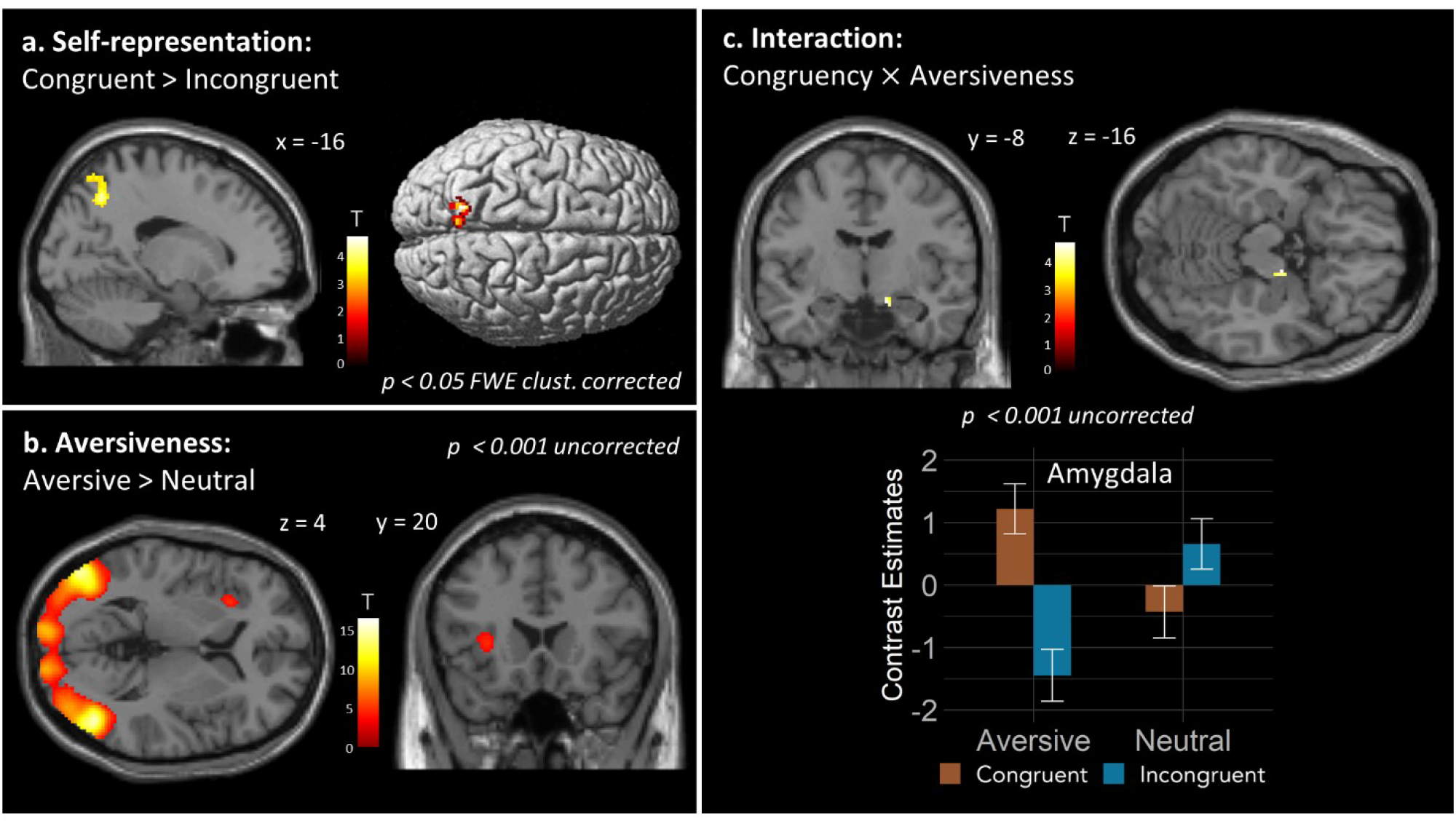
**a**. Congruent versus incongruent visual-tactile stimulation produced significant activation differences in the left SPL (area 7A; *p* < .05, FWE corrected on the cluster level). **b**. Aversive versus neutral stimuli showed significant activation differences in left aIns (area Id7; *p* < .001, uncorrected), and left and right middle temporal area (LOC/hMT+/V5), left and right V1, and right fusiform gyrus (*p* < .05, FWE corrected on the cluster level). **c**. Interaction Congruency × Aversiveness revealed activations in right amygdala (area SF; *p* < .001, uncorrected). Here, activations within anatomical masks of the bilateral amygdala and insula (SPM Anatomy Toolbox; ^**40**^) are shown. Mean contrast estimates of peak activations for both regions are plotted; error bars represent standard error.

#### 3.2.2 Aversive vs. Neutral

Contrasting *aversive* (spider) versus *neutral* (car) conditions revealed significantly higher activation in left and right middle temporal area, comprising Lateral Occipital Complex (LOC) and hMT+/V5, as well as left and right V1, and right fusiform gyrus (FWE corrected; see Figure 2B and Table 1).

When testing in the a priori defined ROIs, activation differences were present in the left dorsal aIns (area Id7; *p* < .001, uncorrected; see Figure 2B and Table 1). No activation differences were found in the amygdala and ACC at a significance level > .001.

#### 3.2.3 Interaction Congruency × Aversiveness

The interaction effect of *Congruency × Aversiveness* did not reveal any significant clusters of activation in the whole brain analysis. When testing in the a priori defined ROIs, activation differences were seen in the right amygdala (superficial area [SF]; *p* < .001, uncorrected; see Figure 2C and Table 1). This interaction effect had a Cohen’s d of 1.08 (large effect size). In the congruent condition, amygdala activity was higher in the presence of an aversive stimulus (vs. neutral), but in the incongruent condition, activity was lower in the presence of an aversive stimulus. Moreover, the difference due to congruency was greater for the aversive condition than the neutral condition. The aIns did not show differences in activation at a significance level > .001.

Post-hoc pairwise comparisons revealed higher amygdala activation for aversive-congruent than aversive-incongruent condition (*p* = .002 Bonferroni corrected; *t(29)* = 4.12), as well as for aversive- congruent compared to neutral-congruent (p = .029, Bonferroni corrected; *t(29)* = 3.05). Finally, the amygdala showed higher activity for the incongruent-neutral compared to incongruent-aversive conditions (*p* = .015, Bonferroni corrected; *t*(29) = -3.30).

When checking for habituation of amygdala activity in response to aversive stimuli, we found modulations of activation in the left (*p* <. 001 within ROI, peak *t* = 3.49; peak *z* = 3.36; *k*_*E*_ = 3; peak x = -18, y = -6, z = -14) and right (*p* <. 001 within ROI, peak *t* = 4.16; peak *z* = 3.96; *k*_*E*_ = 8; peak x = 24, y = -4, z = -12) amygdala.

#### 3.2.4 Control Analyses

Finally, we conducted control analyses to investigate potential effects of individual differences in fear of spiders. Notably, adding individual FSQ scores as a covariate to the 2 × 2 ANOVA with factors Congruency and Aversiveness did not change the results. Particularly, in the interaction Congruency × Aversiveness, a significant activation in the right amygdala was still present with the same activation profile (i.e., contrast estimates) for each pair of conditions, though with a slightly smaller t-value compared to our main analysis (*p* = .001, uncorrected within ROI; peak *t* = 3.19; peak *z* = 3.07; *k*_*E*_ = 3; peak x = 14, y = -6, z = -14). Furthermore, in the 2 × 2 ANOVA with factors Aversive-Congruency (aversive-congruent, aversive-incongruent) and Fear Group (high, low), no interaction effects were seen in the amygdala (*p* >.05, uncorrected within ROI). However, left V5/hMT+ showed an activation (*p* = .001 uncorrected; peak *t* = 3.48; peak *z* = 3.09, *k*_*E*_ = 4, peak x = -58, y = -66, z = 12).

## 4 Discussion

In this study, we investigated brain activity of participants experiencing visual stimulation on a VR arm synchronized with tactile stimulation of the real arm. The aversiveness of the visual stimuli was manipulated, as well as the congruency of the visual and tactile stimuli. The goal was to explore the interplay of emotional processes and self-related multisensory integration.

In response to the retrospective questions regarding body ownership, participants reported experiencing more body ownership during congruent versus incongruent trials. The aversive stimulus (spider) was rated as significantly more “uncomfortable” and “scary”, and less “pleasant” than the neutral stimulus (car). Participants could also clearly discriminate between congruent and incongruent stimulation, and the tactile stimulation was experienced as temporally synchronized with the movement of the visual stimuli. In addition, the results of the control task also revealed that participants were able to focus their attention - as indexed by d’, consistently throughout all runs.

The fMRI results showed higher PPC activity during congruent (vs. incongruent) visuo-tactile stimulation, but neither in the EBA nor the PMv at a significance threshold of p < .001. Additionally, the aIns and visual areas showed higher activation during aversive (vs. neutral) visual stimulation, though neither the amygdala nor the ACC at a significance threshold of p < .001. Finally, testing for interaction effects of Congruency × Aversiveness revealed higher amygdala and pregenual ACC activity during congruent and aversive trials, suggesting that the activation of the amygdala while viewing aversive stimuli depended on the success of the multisensory integration—and embodiment of the artificial limb.

Concerning the manipulation of congruency, the area 7A, corresponding to the posterior SPL, showed significantly higher activation when the direction of the tactile stimulation was congruent (vs. incongruent) with the direction of the visual movement. This result is in line with previous research showing that the posterior SPL is associated with visuo-tactile integration and encoding the internal representation of the body ^44–46^. This suggest that the virtual arm was more integrated into the participant’s own body representation during congruent visuo-tactile stimulation. However, we did not find a significant difference of activation in the EBA between congruent and incongruent conditions. This could be due to the different set-ups between our experiment and previous RHI studies. In particular, in classical RHI paradigms, the fake arm is displaced from the real arm (e.g., ^47^), whereas in the current experiment, the VR arm was at the same visual location as the participant’s real arm. In the former, this displacement creates a visuo-proprioceptive conflict, which has been linked to the activation of the EBA. The EBA is thought to be involved in the integration of the fake arm in the brain’s internal visual body representation and its activity might largely reflect the process of minimizing the prediction error related to conflicting sensory (visual and proprioceptive) signals ^11,47^. In the latter, there was potentially less visuo-proprioceptive conflict, thus no strong involvement of the EBA. Earlier studies have also revealed activation of the PMv, related to multisensory integration and preparation for action ^15,16^. In our study, we did not find significant differences in PMv activity between the congruent and incongruent conditions. This could be due to the relatively short trial duration (2s), the visuo-tactile stimulation ending before the full onset of the RHI. Indeed, previous studies investigating the RHI typically applied stimulation for longer period of time (i.e., 30-35 seconds), and participants reported the start of the illusion 6 to 10 seconds after beginning stimulation ^48,47^. In that context, Ehrsson and colleagues (2004) found that the PMv activity was associated with the after-onset period of the RHI (i.e., approx. 11s after the start of the stroking). Taken together, the questionnaire and these fMRI and results indicate that during the congruent condition, visual and tactile stimuli were more integrated than during the incongruent condition, consistent with previous studies of multisensory integration in the context of body ownership.

Concerning the manipulation of aversiveness, the aIns, which was previously linked to emotional processing ^18,49^, showed higher activation during aversive (vs. neutral) visual stimulation. The aIns is thought to be a hub where the multiple sensory inputs, affective/motivational signals, and visceral information, converge and are integrated in order to detect salient stimuli ^50^. More specifically, the area Id7 belongs to the dorsal part of the aIns ^51^. This area is thought to be involved in various functions, including processing of logic (negation) ^51^, integration of sensory, emotional and cognitive information ^52^, interoception ^53^. Together with the dorsal ACC and amygdala, the dorsal aIns is also part of the salience network ^54^. While ACC appeared in our Aversiveness contrast at a lower threshold, there was no significant difference of activation in the amygdala between aversive and neutral conditions. The activation of dorsal aIns could be due to a difference of saliency between the aversive and neutral stimuli: the aversive stimulus could have been detected as salient, triggering an attentional reorienting in order to facilitate its processing ^55^. This interpretation would also be in line with the activations found in visual areas. The bilateral middle temporal area, comprising LOC and hMT+/V5, the fusiform gyrus and V1 showed significantly higher activations for the aversive conditions than for the neutral conditions. It has been shown that higher visual areas comprising LOC and V5 are more activated for aversive than neutral visual stimuli ^56^, even when controlling for non-emotional potential confounds ^57^ (i.e., colors, visual complexity) or accounting for basic visual perception effects ^58^ (i.e., face and scene perception). Moreover, the middle temporal and fusiform gyri were more activated in spider-phobic participants than in controls when viewing pictures of spider ^59,60^. However, although the two visual stimuli were designed to have identical movement characteristics (i.e., starting and finishing points, distance, speed), we cannot exclude that this higher activation during aversive trials may have resulted from the difference in quality of movement of the stimuli (see Limitations below for an additional control experiment to address this point). Particularly, hMT+/V5 is thought to process visual and tactile motion direction ^61–63^, and whereas the car moved along the arm without changing its shape, the spider’s legs moved to simulate crawling. In addition to that, the aversive and neutral stimuli were not perfectly matched in terms of low-visual features (i.e. colors, shapes). Therefore, one possibility is that the activation in early visual regions (V1) found in the current study could be rather due to a difference in visual features between the two stimuli, and the activation in higher visual areas (fusiform gyrus, LOC, hMT+/V5) could be purely due to the difference of aversiveness. Finally, our control analyses did not show significant effects of FSQ score in the amygdala, though effects were seen in V5/hMT+. It has been shown previously that amygdala activation, in response to viewing pictures of spiders, was correlated to FSQ scores in phobic patients ^64^. In our non-phobic sample, it seems that, while the activity in V5/hMT+ was modulated by the different levels of spider fear, the activity in the amygdala was not.

The main aim of the study was to investigate the interplay between emotional processing and multi-sensory integration, thus the Congruency × Aversiveness effect. The choice of the stimuli in the present experiment was based on a previous study, which found activation of the amygdala when using a video of a spider as a (phylogenetic) threat stimuli with non-phobic participants ^65^. Contrary to what we expected, the amygdala showed no higher activation during aversive trials (versus neutral). Amygdala activity has been shown to be modulated by novelty of the emotional stimuli ^66^ and to decrease when participants are repeatedly exposed to spiders ^65,67^. The results from our control analysis pointed towards a habituation effect during aversive trials, but further studies designed specifically to investigate this phenomenon are needed. Importantly however, the contrast for the interaction between congruency and aversiveness revealed a significant effect on amygdala activity. In the congruent condition, there was higher activation for the aversive compared to the neutral stimuli, but in the incongruent condition this was reversed. The interaction appeared mostly driven by the difference between congruent and incongruent trials in the aversive condition. Surprisingly, the amygdala was less responsive to aversive incongruent than to neutral conditions. The pattern of the interaction suggests that the effect of aversiveness depended on the strength of visual-tactile integration. This pattern could be linked to threat detection and selective attention ^68^. The modulation of attention and the increased response in the visual regions due to emotional stimuli is thought to be modulated by the amygdala ^69^. Previous research has shown that evolutionarily fear-relevant stimuli (including spiders) were detected more quickly (vs. neutral) among distractor stimuli ^70^, and that the amygdala might mediate the capturing of attention when a threat is detected ^71^. Indeed, in healthy participants, attentional blink (i.e., an impairment in the detection of a target if another stimulus precedes it too closely in time) is reduced in the presence of aversive stimuli (vs. neutral), but not in patients with bilateral damage to the amygdala ^72^, indicating that the amygdala plays an important role in the affective modulation of perceptual sensitivity. Therefore, in the context of our study, one possible interpretation could be that, during visuo-tactile congruent trials, the spider may have captured participants’ attention and enhanced perception of the aversive stimulus when the VR arm was perceived more strongly as part of their body, that is, when the stimulus represented a more relevant threat to the bodily self ^73^. The finding of a reduced amygdala activity for the aversive compared to the neutral stimulus in the incongruent condition is puzzling, and more research is needed to understand this effect. Finally, as the ratings for body ownership were not assessed separately for each aversiveness condition, it is possible that the strength of body ownership was either higher or lower during aversive trials than neutral trials. That is, amygdala activation might have been due to the aversiveness that the spider elicited by threatening the arm which was believed to be part of the participant’s body. Alternatively, it is possible that the virtual arm was less incorporated when the spider was seen on it (compared to the toy car), in an attempt to distance the aversive stimulus from the body. In this case, amygdala activation may be rather due to a higher aversive reaction to incorporating the arm into one’s body schema when there is a stimulus threatening it. In summary, we found that amygdala activity in response to an affective stimulus is influenced by the strength of multisensory integration underlying body ownership. This may suggest enhanced perception of aversive stimuli when they represent a bodily threat.

### 4.1 Limitations

One limitation of our study was that the aversive and neutral stimuli were not perfectly matched in terms of low-level visual properties (e.g., colors, shapes), motion type (e.g., biomotion vs. rigid), realism (e.g., possible vs. not possible), familiarity (e.g., more vs. less), visual complexity (e.g. high vs. low), visual and tactile movement consistency (e.g., discrete vibration and discrete footsteps vs. discrete vibration on constant pressure from wheels), and animacy (e.g., agent vs. non agent). This limitation makes it difficult to distinguish between stimulus-versus emotion-driven differences in brain responses. However, in more ecologically valid settings, identical stimuli only differing in emotional loading are almost never found. Our stimuli were matched in terms of size, moving speed, and starting/end point. The stimulus material could nevertheless be improved by generating for example a variety of aversive and neutral stimuli with balanced low-level, non-emotional features. To address some of these possible confounds, we created new aversive and neutral stimuli and ran a visual control experiment (*N* = 23). The new aversive stimuli comprised four spiders only varying in color (brown, red, black, gray), and the neutral stimuli consisted of four non-aversive insects (eight-legged “ladybugs”) in the same colors as the spiders. To perfectly match the amount of motion in the stimuli, both types of stimuli had the same leg movements and number of legs, as well as similar body-shapes. It should be noted, however, that some participants mentioned perceiving the ladybugs as “unnatural” because of the number of legs. Contrasting aversive vs. neutral conditions in this control experiment revealed similar results as found in the current analyses, although voxel clusters were more spatially constrained. Namely, we found significantly higher activation in left and right middle temporal area (comprising LOC and hMT+/V5), as well as left and right V1 and right V3 (*p* < .05 FWE-corrected; see Supplementary Materials for more details). These results suggest that the current findings are not driven solely by visual differences, but indeed by different emotional processing of aversive and neutral stimuli. Nevertheless, the interpretation of the aversiveness contrast remains somehow inconclusive until the effect is replicated with additional stimuli which can control for other potential visual differences between conditions. Further investigation should also seek to elucidate the mechanisms underlying the interaction between aversiveness and body ownership.

A further limitation was that behavioral ratings (e.g., congruency, body ownership, aversiveness) were assessed only retrospectively and could therefore have been influenced by expectations and memory biases. Future studies could avoid this problem by collecting ratings after each individual trial.

Finally, future studies could make the VR experience even more immersive by improving the tactile stimulation and incorporating electrical pulses that more closely mimic the movement of the visual stimuli, therefore potentially increasing both the emotional arousal ^27^ and the body ownership illusion.

## 4.2 Conclusion

Using a novel, fully automated VR-fMRI setup, the interaction between emotion and multisensory integration underlying body ownership was investigated. The findings from this study add to the evidence that the PPC is recruited during visuo-tactile integration and that the aIns is related to aversiveness. More importantly, we found an interaction effect of Congruency × Aversiveness in the amygdala. This new finding points towards an enhanced processing of aversive stimuli when body-related information is more strongly integrated into the bodily representation of the self. Overall, the results show that, with the help of sophisticated VR-fMRI paradigms, important scientific questions can be addressed in a novel way, but that the complexity of the setup also poses new challenges in the interpretation of these findings.

## Data availability

The dataset generated and analyzed during the current study is available from the corresponding author on reasonable request.

## 5 Acknowledgements (optional)

We thank Sarah Polk for English language editing and proof-reading the manuscript.

## 6 Author contributions (names must be given as initials)

AF – Conceptualization, Methodology, Data collection, Data analysis, Writing – original draft, figures, review & editing

TTS – Supervision, Conceptualization, Data collection, Writing – review & editing

TN – Methodology, Data collection, Writing – review & editing

FB – Supervision, Conceptualization, Writing – review & editing, Funding acquisition

## 7 Additional Information (including a Competing Interests Statement)

### Link to the 3D assets used for this study

- **Spider:** https://assetstore.unity.com/packages/3d/characters/creatures/free-fantasy-spider-10104
- **Car:** https://assetstore.unity.com/packages/3d/vehicles/land/retro-cartoon-cars-cicada-96158
- **Arm:** https://assetstore.unity.com/packages/3d/characters/humanoids/vr-hands-and-fp-arms-pack-77815
- **Room:** https://assetstore.unity.com/packages/3d/environments/morgue-room-pbr-65817

## References

1. Macaluso, E. & Driver, J. Multisensory spatial interactions: A window onto functional integration in the human brain. Trends in Neurosciences vol. 28 (2005).

2. Blanke, O. Multisensory brain mechanisms of bodily self-consciousness. Nature Reviews Neuroscience vol. 13 (2012).

3. Rohe, T. & Noppeney, U. Cortical Hierarchies Perform Bayesian Causal Inference in Multisensory Perception. PLoS Biol. 13, (2015).

4. Kilteni, K., Maselli, A., Kording, K. P. & Slater, M. Over my fake body: body ownership illusions for studying the multisensory basis of own-body perception. Front. Hum. Neurosci. (2015) doi:10.3389/fnhum.2015.00141.

5. Samad, M., Chung, A. J. & Shams, L. Perception of body ownership is driven by Bayesian sensory inference. PLoS ONE (2015) doi:10.1371/journal.pone.0117178.

6. Botvinick, M. & Cohen, J. Rubber hands ‘feel’ touch that eyes see [8]. Nature (1998) doi:10.1038/35784.

7. Ehrsson, H. H. The Concept of Body Ownership and Its Relation to Multisensory Integration. In The New Handbook of Multisensory Processes (2012).

8. Tsakiris, M. & Haggard, P. The rubber hand illusion revisited: Visuotactile integration and self-attribution. J. Exp. Psychol. Hum. Percept. Perform. (2005) doi:10.1037/0096-1523.31.1.80.

9. Bekrater-Bodmann, R. et al. The importance of synchrony and temporal order of visual and tactile input for illusory limb ownership experiences - An fMRI study applying virtual reality. PLoS ONE (2014) doi:10.1371/journal.pone.0087013.

10. Ehrsson, H. H., Spence, C. & Passingham, R. E. That’s my hand! Activity in premotor cortex reflects feeling of ownership of a limb. Science 305, 875–877 (2004).

11. Limanowski, J. & Blankenburg, F. Network activity underlying the illusory self-attribution of a dummy arm. Hum. Brain Mapp. (2015) doi:10.1002/hbm.22770.

12. Brozzoli, C., Gentile, G. & Ehrsson, H. H. That’s Near My Hand! Parietal and Premotor Coding of Hand-Centered Space Contributes to Localization and Self-Attribution of the Hand. J. Neurosci. (2012) doi:10.1523/JNEUROSCI.2660-12.2012.

13. Heed, T., Backhaus, J. & Röder, B. Integration of hand and finger location in external spatial coordinates for tactile localization. J. Exp. Psychol. Hum. Percept. Perform. 38, 386–401 (2012).

14. Heed, T., Buchholz, V. N., Engel, A. K. & Röder, B. Tactile remapping: from coordinate transformation to integration in sensorimotor processing. Trends Cogn. Sci. 19, 251–258 (2015).

16. Barany, D. A., Della-Maggiore, V., Viswanathan, S., Cieslak, M. & Grafton, S. T. Feature Interactions Enable Decoding of Sensorimotor Transformations for Goal-Directed Movement. J. Neurosci. (2014) doi:10.1523/JNEUROSCI.5173-13.2014.

17. Craig, A. D. How do you feel? Interoception: The sense of the physiological condition of the body. Nat. Rev. Neurosci. 3, 655–666 (2002).

18. Medford, N. & Critchley, H. D. Conjoint activity of anterior insular and anterior cingulate cortex: Awareness and response. Brain Struct. Funct. 214, 535–549 (2010).

19. Ehrsson, H. H., Wiech, K., Weiskopf, N., Dolan, R. J. & Passingham, R. E. Threatening a rubber hand that you feel is yours elicits a cortical anxiety response. Proc. Natl. Acad. Sci. U. S. A. 104, 9828–9833 (2007).

20. LeDoux, J. The amygdala. Current Biology (2007) doi:10.1016/j.cub.2007.08.005.

21. Peelen, M. V., Atkinson, A. P., Andersson, F. & Vuilleumier, P. Emotional modulation of body-selective visual areas. Soc. Cogn. Affect. Neurosci. 2, 274–283 (2007).

22. Lenggenhager, B., Tadi, T., Metzinger, T. & Blanke, O. Video ergo sum: Manipulating bodily self-consciousness. Science 317, 1096–1099 (2007).

23. Maselli, A., Kilteni, K., López-Moliner, J. & Slater, M. The sense of body ownership relaxes temporal constraints for multisensory integration. Sci. Rep. 6, (2016).

24. Mueller, C. et al. Building virtual reality fMRI paradigms: A framework for presenting immersive virtual environments. J. Neurosci. Methods 209, 290–298 (2012).

25. Reggente, N. et al. Enhancing the ecological validity of fMRI memory research using virtual reality. Frontiers in Neuroscience vol. 12 (2018).

26. Riva, G. et al. Affective interactions using virtual reality: The link between presence and emotions. Cyberpsychol. Behav. 10, 45–56 (2007).

27. Price, M., Mehta, N., Tone, E. B. & Anderson, P. L. Does engagement with exposure yield better outcomes? Components of presence as a predictor of treatment response for virtual reality exposure therapy for social phobia. J. Anxiety Disord. 25, 763–770 (2011).

28. Diemer, J., Alpers, G. W., Peperkorn, H. M., Shiban, Y. & Mühlberger, A. The impact of perception and presence on emotional reactions: a review of research in virtual reality. Front. Psychol. 6, (2015).

29. Gaebler, M. et al. Stereoscopic depth increases intersubject correlations of brain networks. NeuroImage 100, 427–434 (2014).

30. Lorenzetti, V. et al. Emotion regulation using virtual environments and real-time fMRI neurofeedback. Front. Neurol. 9, (2018).

31. Davey, G. C. L. Characteristics of individuals with fear of spiders. Anxiety Res. 4, 299–314 (1991).

32. Oldfield, R. C. The assessment and analysis of handedness: The Edinburgh inventory. Neuropsychologia 9, 97–113 (1971).

33. Macey, P. M., Macey, K. E., Kumar, R. & Harper, R. M. A method for removal of global effects from fMRI time series. NeuroImage 22, 360–366 (2004).

34. Kleiner, M. et al. What’s new in psychtoolbox-3. Perception 36, 1–16 (2007).

35. Dodgson, N. A. Variation and extrema of human interpupillary distance. in Stereoscopic Displays and Virtual Reality Systems XI vol. 5291 36–46 (International Society for Optics and Photonics, 2004).

36. Ashburner, J. A fast diffeomorphic image registration algorithm. NeuroImage 38, 95–113 (2007).

37. Behzadi, Y., Restom, K., Liau, J. & Liu, T. T. A component based noise correction method (CompCor) for BOLD and perfusion based fMRI. NeuroImage 37, 90–101 (2007).

38. Macmillan, N. A. & Kaplan, H. L. Detection Theory Analysis of Group Data. Estimating Sensitivity From Average Hit and False-Alarm Rates. Psychol. Bull. 98, 185–199 (1985).

39. Stanislaw, H. & Todorov, N. Calculation of signal detection theory measures. Behav. Res. Methods Instrum. Comput. 31, 137–149 (1999).

40. Eickhoff, S. B. et al. A new SPM toolbox for combining probabilistic cytoarchitectonic maps and functional imaging data. NeuroImage 25, 1325–1335 (2005).

41. Moayedi, M. et al. The structural and functional connectivity neural underpinnings of body image. Hum. Brain Mapp. 42, 3608–3619 (2021).

42. Szymanski, J. & O’Donohue, W. Fear of Spiders Questionnaire. J. Behav. Ther. Exp. Psychiatry 26, 31–34 (1995).

43. de Haan, A. M., Smit, M., Van der Stigchel, S. & Dijkerman, H. C. Approaching threat modulates visuotactile interactions in peripersonal space. Exp. Brain Res. 234, 1875–1884 (2016).

44. Wolpert, D. M., Goodbody, S. J. & Husain, M. Maintaining internal representations: The role of the human superior parietal lobe. Nat. Neurosci. 1, 529–533 (1998).

45. Graziano, M. & Botvinick, M. M. How the brain represents the body: Insights from neurophysiology and psychology. Atten. Perform. Vol XIX Common Mech. Percept. Action 136–157 (2002).

46. Limanowski, J. & Blankenburg, F. Integration of visual and proprioceptive limb position information in human posterior parietal, premotor, and extrastriate cortex. J. Neurosci. 36, 2582–2589 (2016).

47. Limanowski, J., Lutti, A. & Blankenburg, F. The extrastriate body area is involved in illusory limb ownership. NeuroImage 86, 514–524 (2014).

48. Petkova, V. I. et al. From part-to whole-body ownership in the multisensory brain. Curr. Biol. 21, 1118–1122 (2011).

49. Craig, A. D. How do you feel - now? The anterior insula and human awareness. Nature Reviews Neuroscience vol. 10 (2009).

50. Menon, V. & Uddin, L. Q. Saliency, switching, attention and control: a network model of insula function. Brain Struct. Funct. 214, 655–667 (2010).

51. Grodzinsky, Y. et al. Logical negation mapped onto the brain. Brain Struct. Funct. 225, 19–31 (2020).

52. Simmons, W. K. et al. Keeping the body in mind: Insula functional organization and functional connectivity integrate interoceptive, exteroceptive, and emotional awareness. Hum. Brain Mapp. 34, 2944–2958 (2013).

53. Kurth, F., Zilles, K., Fox, P. T., Laird, A. R. & Eickhoff, S. B. A link between the systems: functional differentiation and integration within the human insula revealed by meta-analysis. Brain Struct. Funct. 214, 519–534 (2010).

54. Seeley, W. W. The Salience Network: A Neural System for Perceiving and Responding to Homeostatic Demands. J. Neurosci. Off. J. Soc. Neurosci. 39, 9878–9882 (2019).

55. Menon, V. Salience Network. in Brain Mapping: An Encyclopedic Reference vol. 2 597–611 (2015).

56. Garrett, A. S. & Maddock, R. J. Separating subjective emotion from the perception of emotion-inducing stimuli: An fMRI study. NeuroImage 33, 263–274 (2006).

57. Taylor, S. F., Liberzon, I. & Koeppe, R. A. The effect of graded aversive stimuli on limbic and visual activation. Neuropsychologia 38, 1415–1425 (2000).

58. Sabatinelli, D. et al. Emotional perception: Meta-analyses of face and natural scene processing. NeuroImage 54, 2524–2533 (2011).

59. Dilger, S. et al. Brain activation to phobia-related pictures in spider phobic humans: An event-related functional magnetic resonance imaging study. Neurosci. Lett. 348, 29–32 (2003).

60. Schienle, A., Schäfer, A., Walter, B., Stark, R. & Vaitl, D. Brain activation of spider phobics towards disorder-relevant, generally disgust-and fear-inducing pictures. Neurosci. Lett. 388, 1–6 (2005).

61. Grossman, E. et al. Brain areas involved in perception of biological motion. J. Cogn. Neurosci. 12, 711–720 (2000).

62. Hagen, M. C. et al. Tactile motion activates the human middle temporal/V5 (MT/V5) complex. Eur. J. Neurosci. 16, 957–964 (2002).

63. Van Kemenade, B. M. et al. Tactile and visual motion direction processing in hMT+/V5. NeuroImage 84, 420–427 (2014).

64. Caseras, X. et al. Dynamics of brain responses to phobic-related stimulation in specific phobia subtypes. Eur. J. Neurosci. 32, 1414–1422 (2010).

65. Mobbs, D. et al. Neural activity associated with monitoring the oscillating threat value of a tarantula. Proc. Natl. Acad. Sci. U. S. A. 107, 20582–20586 (2010).

66. Weierich, M. R., Wright, C. I., Negreira, A., Dickerson, B. C. & Barrett, L. F. Novelty as a dimension in the affective brain. NeuroImage 49, 2871–2878 (2010).

67. Björkstrand, J. et al. Decrease in amygdala activity during repeated exposure to spider images predicts avoidance behavior in spider fearful individuals. Transl. Psychiatry 10, 1–10 (2020).

68. Bishop, S. J. Neural Mechanisms Underlying Selective Attention to Threat. Ann. N. Y. Acad. Sci. 1129, 141–152 (2008).

69. Vuilleumier, P. How brains beware: neural mechanisms of emotional attention. Trends Cogn. Sci. 9, 585–594 (2005).

70. Öhman, A., Flykt, A. & Esteves, F. Emotion drives attention: Detecting the snake in the grass. J. Exp. Psychol. Gen. 130, 466–478 (2001).

71. Öhman, A. The role of the amygdala in human fear: Automatic detection of threat. Psychoneuroendocrinology 30, 953–958 (2005).

72. Anderson, A. K. & Phelps, E. A. Lesions of the human amygdala impair enhanced perception of emotionally salient events. Nature 411, 305–309 (2001).

73. Sander, D., Grafman, J. & Zalla, T. The human amygdala: an evolved system for relevance detection. Rev. Neurosci. 14, 303–316 (2003).

74. Zimmermann, M., Meulenbroek, R. G. J. & De Lange, F. P. Motor planning is facilitated by adopting an action’s goal posture: An fMRI study. Cereb. Cortex 22, 122–131 (2012).

