## Supplementary Materials for "Enhanced processing of aversive stimuli on embodied artificial limbs by the human amygdala"

**Supplementary Figure S1.** Behavioral ratings from the post-scan questionnaire

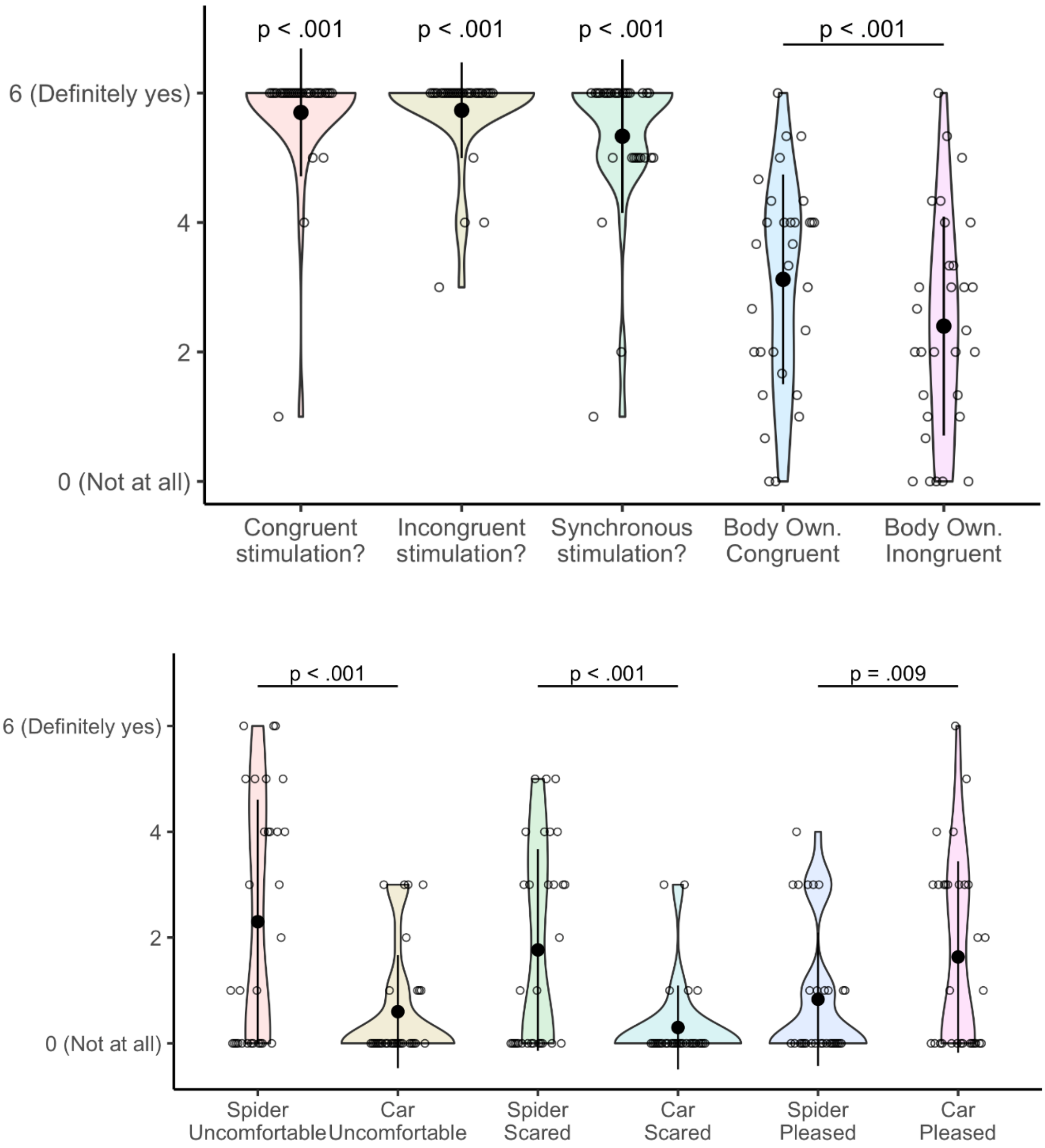

**Supplementary Table S2.** Descriptive statistics of the behavioral ratings from the post-scan questionnaire.

**Table S2**

*Descriptive statistics and results of Wilcoxon tests (against 0 for the validation items, comparing across conditions for body ownership scores and affectivity ratings) of numerical rating scales where 0 = "Not at all" and 6 = "Definitely yes".*

| Validation items | | M (SD) | $Z$ | $p$ |
|---|---|---|---|---|
| Was the tactile stimulation for some trials going the same direction as the visual moving object? | | 5.70 (0.99) | 5.12 | < .001 |
| Was the tactile stimulation for some trials going the opposite direction as the visual moving object? | | 5.73 (0.74) | 5.15 | < .001 |
| Was the visual moving object synchronized to the tactile stimulation? | | 5.33 (1.18) | 4.73 | < .001 |
| | Congruent | Incongruent | $Z$ | $p$ |
| Body ownership scores | 3.12 (1.62) | 2.40 (1.69) | 3.27 | < .001 |
| Did the moving object make you feel... | Spider | Car | $Z$ | $p_{FDR}$ |
| uncomfortable? | 2.3 (2.31) | 0.6 (1.07) | 3.72 | < .001 |
| scared? | 1.8 (1.91) | 0.3 (0.79) | 3.51 | < .001 |
| pleased? | 0.8 (1.26) | 1.6 (1.81) | 2.56 | .009 |

*Note.* M = mean, SD = standard deviation.

Body ownership scores were calculated as the average rating across the following questions: "I felt as I was looking at my own arm and hand," "I felt as if the virtual arm and hand was part of my body," and "I felt as if the virtual arm and hand were my arm and hand."

**Supplementary Figure S3.** Visual control experiment. **a.** The aversive stimuli comprised four spiders only varying in color (brown, red, black, gray), and the neutral stimuli comprised four non-aversive insects (i.e., eight-legged "ladybugs") in the same colors as the spiders. Note that the two types of stimuli had similar body-shapes and the same number of legs and leg movements. **b.** Aversive versus neutral stimuli showed significant activation differences in left and right middle temporal area (LOC/hMT+/V5), left and right V1, and V3 ($p < .05$, FWE corrected on the cluster level). There were no significant differences in left aIns.

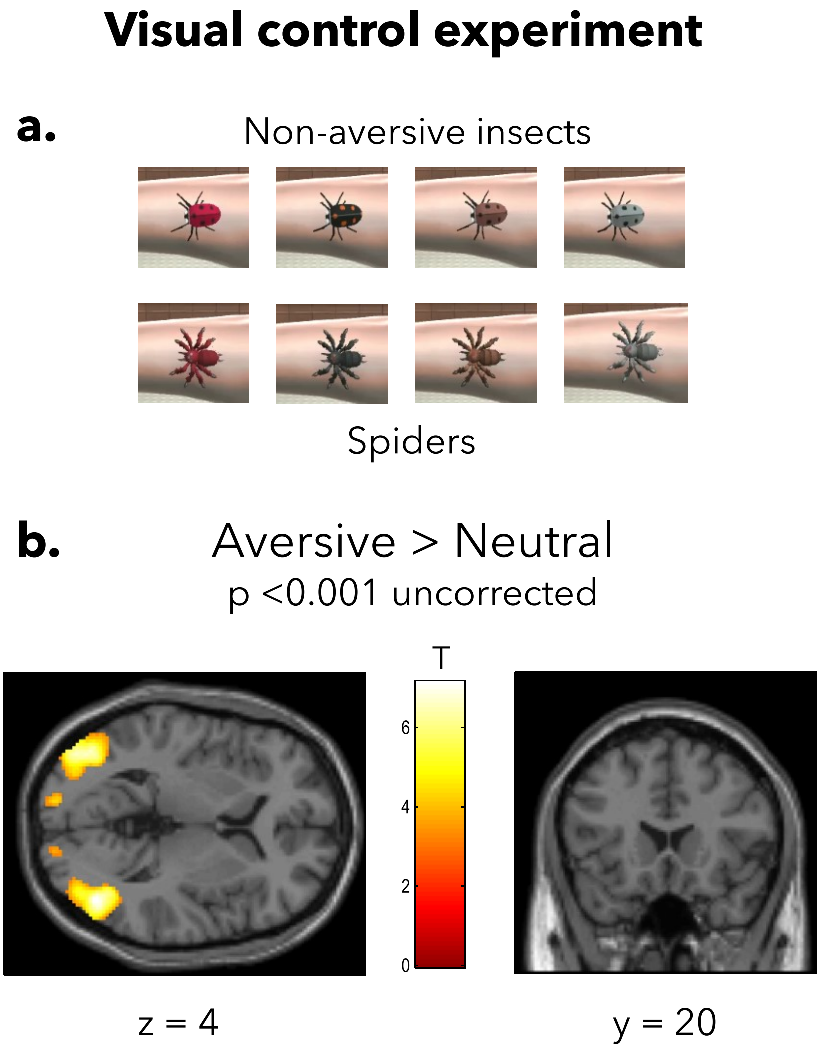
